# Mixture-models for stimulus-selective stopping

**DOI:** 10.1101/2025.11.14.688422

**Authors:** Paria Jahansa, Adele Diederich, Hans Colonius

## Abstract

Stimulus-selective stopping extends the standard stop signal task by occasionally presenting an ‘ignore’ signal instead of a stop signal, in which case participants are instructed to continue responding to the go signal. Here we present several model classes that are based on the idea that responses observed under an ignore signal are the result of a probabilistic mixture from the processing distributions of the go and the ignore signal. Earlier work ensures that the mixture hypothesis is statistically testable. We derive quantitative predictions and parameter estimation for model classes that differ in the way the mixture is introduced. The results are illustrated with an application to a published dataset for stimulus-selective stopping.

## Stop-signal paradigm and selective inhibition

In the stop-signal paradigm, participants engage in a reaction time task where they respond by pressing a button upon encountering a *go* signal. Occasionally, a *stop* signal is presented after a variable stop-signal delay (SSD), prompting participants to withhold their response. When SSD is long, participants typically fail to inhibit their response, resulting in a recorded reaction time (RT). Conversely, for short SSDs, participants often succeed in following the instruction, leading to no recorded response. This paradigm is widely used for investigating response inhibition in both simple and choice reaction time tasks (e.g., Matzke, Verbruggen, & Logan, 2018). When a subject succeeds in canceling the response in the presence of a stop signal, no RT can be recorded. Thus, a main goal of modeling performance in this task is to somehow obtain an estimate of the non-observable time to process the signal. In the prevalent model, known as the *race model* (Logan & Cowan, 1984), performance in the stop-signal task is represented as a race between two stochastically independent random variables representing the go and stop signal processing times, denoted as *T*_*go*_ and *T*_*stop*_, respectively. Whenever *T*_*go*_ takes on a value smaller than the one *T*_*stop*_ +*t*_*s*_ takes on (*t*_*s*_ denoting the value of SSD), then a response is given in spite of the stopping instructions. Estimation of the stop signal processing time (aka SSRT), i.e. the non-observable time to process the stop signal, relies on a further assumption of the model known as *context invariance*^1^: *the distribution of T*_*go*_ is the same with and without a stop signal being present in a trial (Colonius, 1990).

More recently, a more complex type of inhibition performance has raised interest, namely *selective inhibition* (Bissett & Logan, 2014). Two main variants exist: in *response-selective* stopping subjects make two responses (usually bimanually) on each trial, and stop signals indicate which response has to be stopped. This has been investigated by an extension of the original race model by Gronau and colleagues (Gronau, Hinder, Salomoni, Matzke, & Heathcote, 2024; Salomoni, Gronau, Heathcote, Matzke, & Hinder, 2023). Our focus here is on the other variant of the stop signal paradigm, known as *stimulus-selective* stopping, where instead of the stop signal sometimes a third type of signal –called *ignore signal* – is presented requiring the participant to continue responding to the go signal. Clearly, the difficulty of this task depends on how easy it is to discriminate between the stop and the ignore signal (see, e.g., Sanchez-Carmona, Rincon-Perez, Lopez-Matin, Albert, & Hinojasa, 2021).

## Strategies in stimulus-selective stopping

In stimulus-selective stopping, one of three, mutually exclusive, stimulus presentations occurs in each trial: (i) only the go signal is presented; (ii) the go signal and, after SSD (*t*_*s*_), a stop signal is presented; (iii) the go signal and, after ISD (*t*_*i*_), an ignore signal is presented^2^. A typical observation is that responses in the presence of an ignore signal are considerably delayed suggesting that stimulus-selective stopping somehow impairs the go process (Sanchez-Carmona et al., 2021; Sanchez-Carmona, Santaniello, Capilla, Hinojasa, & Albert, 2019; van de Laar, van den Wildenberg, van Boxtel. G.J.M., & van der Molen, 2011). Two main alternative explanations for the effect have been discussed up to now. First, subjects may stop whenever a signal, whether stop or ignore, occurs and then restart if the signal is an ignore signal, denoted as the *Stop-then-Discriminate* strategy. Second, subjects may discriminate the signal before deciding to stop. If the signal is a stop signal, they stop; if the signal is an ignore signal, they complete the go process as usual, without ever initiating the stop process; this is the *Discriminate-then-Stop* strategy (for further details, see Bissett & Logan, 2014).

Several studies have explored the nature of differences between strategies or factors that may influence strategy adoption. For example, Bissett and Logan (2014) investigated whether stopping and recognizing signals in a selective stopping task occur independently or interfere with each other. Participants were required to stop their responses when shown a stop signal but continue responding when shown an ignore signal, with varying difficulty in distinguishing between them. The key findings revealed that many participants did not treat stopping and signal discrimination as separate processes. Instead, the need to differentiate between stop and ignore signals interfered with their ability to stop efficiently. When the signals were harder to distinguish, stopping performance decreased significantly, indicating that stopping was affected by the complexity of signal recognition. These results challenge traditional models of selective stopping, which assume independence between stopping and discrimination. Instead, the study suggests that the interaction between these processes plays a crucial role in response control, showing that stopping efficiency is directly influenced by how difficult it is to recognize signals.

Sanchez-Carmona et al. (2021) investigated a stimulus-selective stop-signal task aiming at determining whether the difficulty in discriminating between stop and ignore signals affects strategy adoption. In this task, a new stimulus –either stop or ignore– was presented shortly after the go stimulus on 40% of trials. The SSD and ISD were initially set to 250 ms and were dynamically adjusted by +50 ms or *−*50 ms based on the participant’s previous performance. Participants employed different strategies, with the Discriminate-then-Stop strategy being dominant in easier discrimination conditions and the Stop-then-Discriminate strategy prevailing in more challenging conditions. These findings align with Bissett and Logan (2014), who suggested that strategy adoption may reflect a conservative approach when stopping is difficult. Several factors influencing strategy selection have also been investigated, including genetics (dopaminergic polymorphisms), task-related variables (signal frequency, discrimination difficulty), proactive inhibition, attentional capture, and working memory. This suggests that while participants generally adapt their strategy based on task demands, some individuals consistently adhere to a single strategy.

## Mixture-models of stimulus-selective stopping

The popularity of the classic, non-parametric race model (Logan & Cowan, 1984) is arguably due to the simplicity of its assumptions and the resulting strong testability. Given the increased complexity of the stimulus-selective stopping task, some modification of the classic race model cannot be avoided, however.

There are three experimental contexts to distinguish, denoted as *𝒢𝒪, 𝒮𝒯𝒪𝒫*, and *ℐ𝒢𝒩𝒪ℛℰ*^3^ which are defined by the possible stimulus types presented in a given trial. For each stimulus type, occurrence of a stimulus triggers realization of a random variable representing the processing time for the stimulus: *T*_*go*_, *T*_*stop*_, and *T*_*ig*_, respectively. In the classic race model, observability of *T*_*stop*_ is limited because no responses are registered when the stopping process is the “winner of the race” between *T*_*go*_ and *T*_*stop*_. In the stimulus-selective paradigm, processing time *T*_*ig*_ is also not directly observable because responses of a participant in context *ℐ𝒢𝒩𝒪ℛℰ* may be due to either responding to the go or the ignore signal.

One way to deal with the non-observability of the distributions of stop and ignore signal processing times is to assume that they belong to a (more or less) plausible family of parametric distribution functions, like ex-Gaussian distributions, as used in Gronau et al. (2024) for modeling response-selective stopping. In order to stay as close as possible to the classic Logan-Cowan race model, here we propose a *semi-parametric* approach that is in-between the two. Specifically, we do not assume a particular distribution (family) but allow for numerical parameters relating the distributions.

### Mixture hypothesis

If a second stimulus occurs after the go signal, participant’s response depends on whether a stop signal or an ignore signal is presented. A response in a stop signal trial is given only if its processing terminates after a response to the go signal has been generated, as in the classic race model. A response in an ignore signal trial is triggered either by the go signal or the ignore signal. Note that assuming a race between go and ignore signal is ruled out because it would predict faster reactions in the presence of ignore signals than without ignore signals, in contradiction to the typical observation of delayed responses in the presence of ignore signals (e.g., Bissett & Logan, 2014). Instead, we propose that a participant’s response is based on processing only one of the signals, either the go signal or the ignore signal, that is, the observed distribution of responses is the result of mixing both distributions. Thus, the fundamental assumption of the mixture model is that the observed responses come from a distribution that is a binary mixture of the distributions for go and ignore signal processing times. The probability of the response being elicited from the ignore signal depends on when it is presented: the later it is given, the smaller the probability that the response has been triggered by it (“late-ignore signal hypothesis”, see below).

Writing 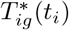 for the random variable representing response time when an ignore signal occurs with a delay *t*_*i*_ (ISD) after the go signal, its distribution function 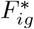 is postulated to be the binary mixture

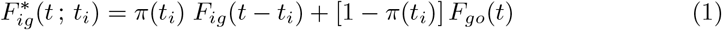

with *F*_*ig*_(*t*) the distribution function of *T*_*ig*_, *π*(*t*_*i*_) the probability that the response is triggered by the ignore signal presented at ISD equal to *t*_*i*_, and *F*_*go*_(*t*) the distribution function of *T*_*go*_. Although the mixture component distribution *F*_*ig*_(*t*) nor the mixture probability *π*(*t*_*i*_) are observable, the mixture hypothesis is nonetheless testable, due to a remarkable result from Falmagne (1968) to be detailed below,

### The general mixture model

Extending the classic independent stop signal model (without ignore signals), we distinguish 3 different trial types, called *context*, according to which stimuli are being presented in a given trial: (i) context *𝒢𝒪* with only a go signal presented, (ii) context *𝒮𝒯𝒪𝒫* where a go signal is followed by a stop signal, and (iii) context *ℐ𝒢𝒩𝒪ℛℰ* where a go signal is followed by an ignore signal. We limit discussion to a simple design: there is 1 go signal, 1 stop signal, and 1 ignore signal, and, typically, the incidence of stop and ignore trials is relatively low and equal.

The general mixture model is defined by the following assumptions:

(A1) in context *𝒢𝒪*: random variable *T*_*go*_ determines the response with distribution function *F*_*go*_(*t*);

(A2) in context *𝒮𝒯𝒪𝒫*: random variable min*{T*_*go*_, *T*_*stop*_ + *t*_*s*_*}* determines the response, like in the classic model (*t*_*s*_ the stop signal delay, SSD)

(A3) in context *ℐ𝒢𝒩𝒪ℛℰ*: the response is determined either by the go signal or the ignore signal; specifically, the distribution of responses is generated by the binary mixture distribution in Equation (1).

(A4) Stochastic independence (SI) holds between *T*_*go*_ and *T*_*stop*_ in context *𝒮𝒯𝒪𝒫* and between *T*_*go*_ and *T*_*ig*_ in context *ℐ𝒢𝒩𝒪ℛℰ*.

(A5) Context invariance holds for *T*_*go*_ across all contexts.

The go signal occurs in all three contexts and, by assuming context invariance (A5), its processing time can be represented by one and the same random variable, *T*_*go*_, in all of them. Because there is no context in which *T*_*stop*_ and *T*_*ig*_ co-occur, no assumption about their stochastic dependency must be made. For simplicity, assumption (A4) states stochastic independence between the variables in contexts *𝒮𝒯𝒪𝒫* and *ℐ𝒢𝒩𝒪ℛℰ*, but that could be modified if data suggest.

Because of the mixture hypothesis (A3), a response observed in context *ℐ𝒢𝒩𝒪ℛℰ* cannot unequivocally be attributed to the subject responding either the go or the ignore signal. If the ignore signal is presented rather late, the response is more likely to have been triggered by the go signal. More specifically, we state this as

#### Late-Ignore-Signal Hypothesis

*In context ℐ𝒢𝒩𝒪ℛε, the probability that the response is triggered by the ignore signal, π*(*t*_*i*_), *is decreasing in t*_*i*_, *the value of ISD*.

### Test of the mixture hypothesis (Falmagne, 1968)

A test of the mixture hypothesis is based on a well-known result:

#### Proposition 1

*(Falmagne, 1968) Let F*_1_ *and F*_2_, *with parameters p*_1_ *and p*_2_ *respectively, be two mixtures of distribution functions H and K. If p*_1_ = *p*_2_, *and if F*_1_(*x*_0_) = *F*_2_(*x*_0_) = *α for some x*_0_, *then any mixture of H and K has value α at x*_0_.

Thus, all binary mixtures of the same two distributions functions intersect at the same point, if they intersect at all. The proposition, referred to as *fixed-point property*, was proved for mixtures of distribution functions. But it also holds for probability density functions, if they exist:

#### Proposition 2

*Let f*_1_ *and f*_2_, *with parameters p*_1_ *and p*_2_ *respectively, be two mixtures of density functions h and k. If p*_1_ ≠ *p*_2_, *and if f*_1_(*x*_0_) = *f*_2_(*x*_0_) = *α for some x*_0_, *then any mixture density of h and k has value α at x*_0_.

*Proof*. By the hypothesis,

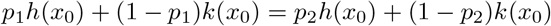

and thus

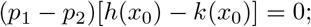

then, since *p*_1_ ≠ *p*_2_, it follows that *h*(*x*_0_) = *k*(*x*_0_); but then,

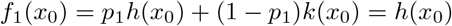

and

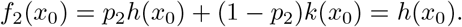

Thus, density mixtures *f*_1_ and *f*_2_ do not depend on *p*_1_, *p*_2_. We conclude that all binary density mixtures of *h* and *k* must intersect at *x*_0_.

The above proof completely parallels the one for Proposition 1 (see also van Maanen, de Jong, & van Rijn, 2014). A few remarks are in order:

1. The point of intersection in Proposition 1 is typically not identical to the one of Proposition 2. For example, if two (mixture) distributions functions are identical up to a shift, there is no point of intersection, whereas the corresponding densities may well intersect at some point.
2. There may exist more than one point of intersection both in case of densities and distribution functions.

Using Falmagne’s result in practice is not that straightforward because one would usually like to “prove” the null hypothesis of the presence of a mixture. A suitable method to compute intersection points and a statistical test to assess the probability that indeed the observed response times are generated by a mixture has been developed by van Maanen and colleagues (van Maanen, Couto, & Lebreton, 2016; van Maanen et al., 2014) (see also Couto, Lebreton, & van Maanen, 2024). Being based on kernel density estimation for histograms, it probes existence of intersection points for mixtures of densities, not distribution functions. This is no limitation, however, because evidence for a mixture of densities implies that the corresponding distribution functions are mixtures as well, although they may not intersect.

### How to test the general mixture model

Testing the mixture hypothesis is a central part of testing the general mixture model. If there is support for the hypothesis, the next step is to probe the race model inequality.

Note that the mixture model is identical to the classic race model in contexts *𝒢𝒪* and *𝒮𝒯𝒪𝒫*. Thus, an important prediction of the latter continues to be valid: the race model inequality under stochastic independence, aka known as the probability inequality test, i.e.

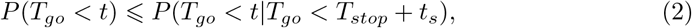

where *t*_*s*_ denotes the stop-signal delay (SSD); for relaxing the assumption see e.g., Colonius, Jahansa, Joe, and Diederich (2023).

However, a further testing of the general mixture model in its non-parametric form is rather limited. Consider Equation (1) again:

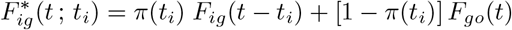

This could be solved for the unknown *F*_*ig*_(*t − t*_*i*_) but is not useful because *π*(*t*_*i*_) is not known, either. Setting *t*_*i*_ “large enough”, Equation (1) reduces to [1 *− π*(*t*_*i*_)] *F*_*go*_(*t*)) for *t > t*_*i*_, thus yielding an estimate of *π*(*t*_*i*_) because *F*_*go*_(*t*) is known by context invariance. Letting *t → ∞*, the general mixture model predicts that the probability to respond in context *ℐ𝒢𝒩𝒪ℛℰ* approaches 1.

Introducing specific distributions for the mixture components would be a possible way to go. For the time being, we prefer a *semi-parametric* approach that allows us to estimate the non-observable parts of the model. Specifically, we assume *F*_*ig*_ to be equal to *F*_*go*_ up to a shift parameter.

## Mixture model with shift

Due to context invariance, *F*_*go*_(*t*) occurs in context *ℐ𝒢𝒩𝒪ℛℰ*, whereas *F*_*ig*_(*t*) is unknown. We now assume that the distribution of *T*_*ig*_ is simply a shifted version of the distribution of *T*_*go*_:

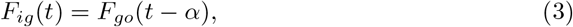

where *α* is a positive constant.

This is the *mixture model with shift*, or MMS model, for short. It postulates that processing of the ignore signal is delayed by a constant *α*, which may be interpreted as the time it takes to discriminate it from a potential stop signal. Although this is a rather strong assumption, it is empirically testable. Moreover, it predicts that the more difficult the discrimination is, the larger *α* should be.

Taking into account the ISD values (*t*_*i*_), the model equation for responses in the *ℐ𝒢𝒩𝒪ℛℰ* context becomes:

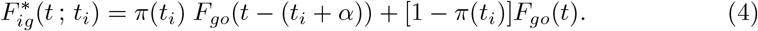

Assuming that an estimate of *F*_*go*_(*t*) is available from context *𝒢𝒪*, testing the model requires estimation of *α* and the mixture probabilities *π*(*t*_*i*_).

### Parameter estimation for the mixture model with shift (MMS)

With *n* different ISD values, there are *n* model equations (4) and *n* mixture probabilities *π*(*t*_*i*_); adding shift parameter *α*, the total number of parameters to be estimated equals *n* + 1.

One way to proceed is taking means (expected values) in (4) yielding (see Appendix)

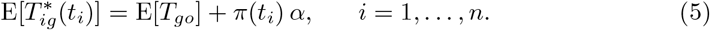

Moreover, in order to estimate *α* we derive a relation between the second (raw) moments of 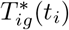 and *T*_*go*_:

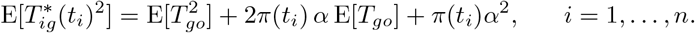

For the derivation, see Appendix. Sample estimates for all (2*n*) expected value terms are available in order to estimate the *n* unknown mixture probabilities plus parameter *α*. Because 2*n > n* + 1 for *n ≥* 2, at least 2 different ISD (*t*_*i*_) are required for testing the model.

## The quantile-mixture model

The idea that ignore signal and go signal distribution only differ by a shift parameter *α*, as expressed in Equation (3), can, arguably, introduced more directly at the level of quantiles. First, we have to introduce the definition of quantile function:

### Definition 1

*Let X be a real-valued continuous random variable with distribution function F* (*x*) *which is continuous from the right. The* quantile function *Q*_*X*_ (*u*) is defined as

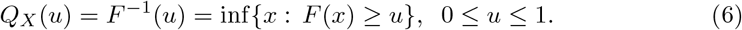

Thus, if there exists an *x* such that *F* (*x*) = *u*, then *F* (*Q*_*X*_ (*u*)) = *u* and *Q*_*X*_*u*) is the smallest value of *x* satisfying *F* (*x*) = *u*. Simplifying the exposition, we will assume that the probability distributions here are continuous and strictly increasing, so that *Q*_*X*_ (*u*) is the unique value of *x* such that *F* (*x*) = *u*. Before introducing the shift assumption, let us briefly consider a more general version of the model.

Let 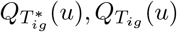 and 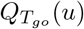 be the quantile functions for the random variables 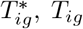, and *T*_*go*_, respectively. The *quantile-mixture model* is identical to the general mixture model defined by assumptions (A1) - (A5), except that the mixture assumption (A3) is replaced by a mixture at the level of quantiles, that is,

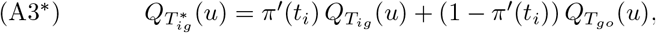

omitting, for simplicity, reference to the *t*_*i*_ value. The primes are used here to distinguish the current parameters from those in the mixture model (MMS) above. Note that quantile function 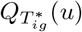, in general differs from the quantile function of the mixture distribution 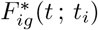. Thus, in principle, data may be consistent with the probability mixture model but not the quantile mixture model, or vice versa^4^. This becomes an important point when parametric distribution assumptions are considered. For a systematic comparison of both model types in a non-parametric reaction time context, we refer to Dixon (2012).

### The quantile-mixture model with shift (QMS)

Given the non-observability of 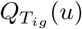, parameter estimates for the quantile mixture model of Equation (3) are available only after further specification. Assuming that the ignore quantile function is simply a shift of the go quantile function

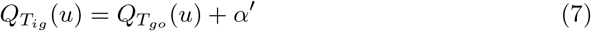

leads to the model equation defining the *quantile-mixture model with shift* (QMS):

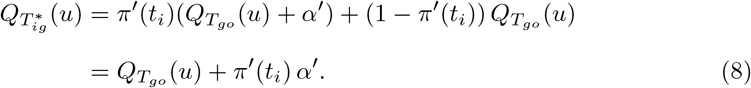

This model equation is similar to the one for the mixture model with shift (MMS), Equation (5),

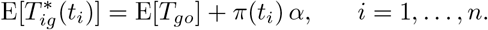

One might even suspect that the latter could be derived from the former, given that

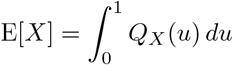

for continuous *X*. Integrating (8) yields

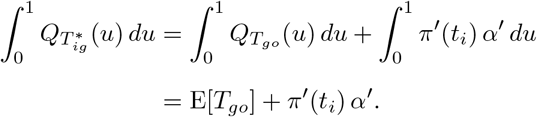

However,

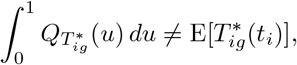

because 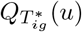 is in general not the quantile function of 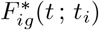, as was explained above.

### Illustrating test of the mixture hypothesis and model parameter estimation with data from Bompas, Sumner, and Hedge (2025)

We chose a manual letter discrimination and selective stopping dataset from Bompas et al. (2025). Bompas and colleagues did not seek to probe specific models of selective stopping but, rather, tried to identify what they called the *non-decision time components* of the response from behavior data alone (for details we must refer to their paper). Our goal here simply is to illustrate the feasibility of testing the mixture hypothesis and of estimating parameters in the models introduced above. Developing procedures for statistically testing and comparing the various models against each other requires further work and will involve an enlargement of empirical datasets.

The dataset includes data from 14 participants; in each block, a triplet of letters was introduced, including two target letters (e.g., Q and O) and one stop letter (e.g., G). Target letters were to be discriminated, pressing the corresponding letter on the keyboard. Stop letters were associated with an instruction to withhold the response. Triplets were chosen such that the two target letters were clearly separated horizontally on the keyboard, and the stop letter sat in between. The second letter was either the same target letter (ignore trials), or the stop letter (stop trials). Ignore and stop trials were presented with equal proportion, with SSDs/ISDs of 83, 100, 117 or 133 ms. The analyses were conducted on 360 single target trials, 240 double letter ignore trials and 240 double letter stop trials in each of 30 blocks.

### Test of the general mixture model: probability inequality

Because, for context *𝒮𝒯𝒪𝒫*, the general mixture model is identical to the classic race model, it is amenable to the usual probability inequality test as stated in Equation (2). Figure 1 shows the sample distribution functions corresponding to the 4 stop signal delays (SSD) compared with the sample go signal distribution. Visual inspection reveals that the inequality is mostly satisfied, except for the shortest SSD at higher response times. In sum, no strong evidence against the model is found from this test.

**Fig. 1:**
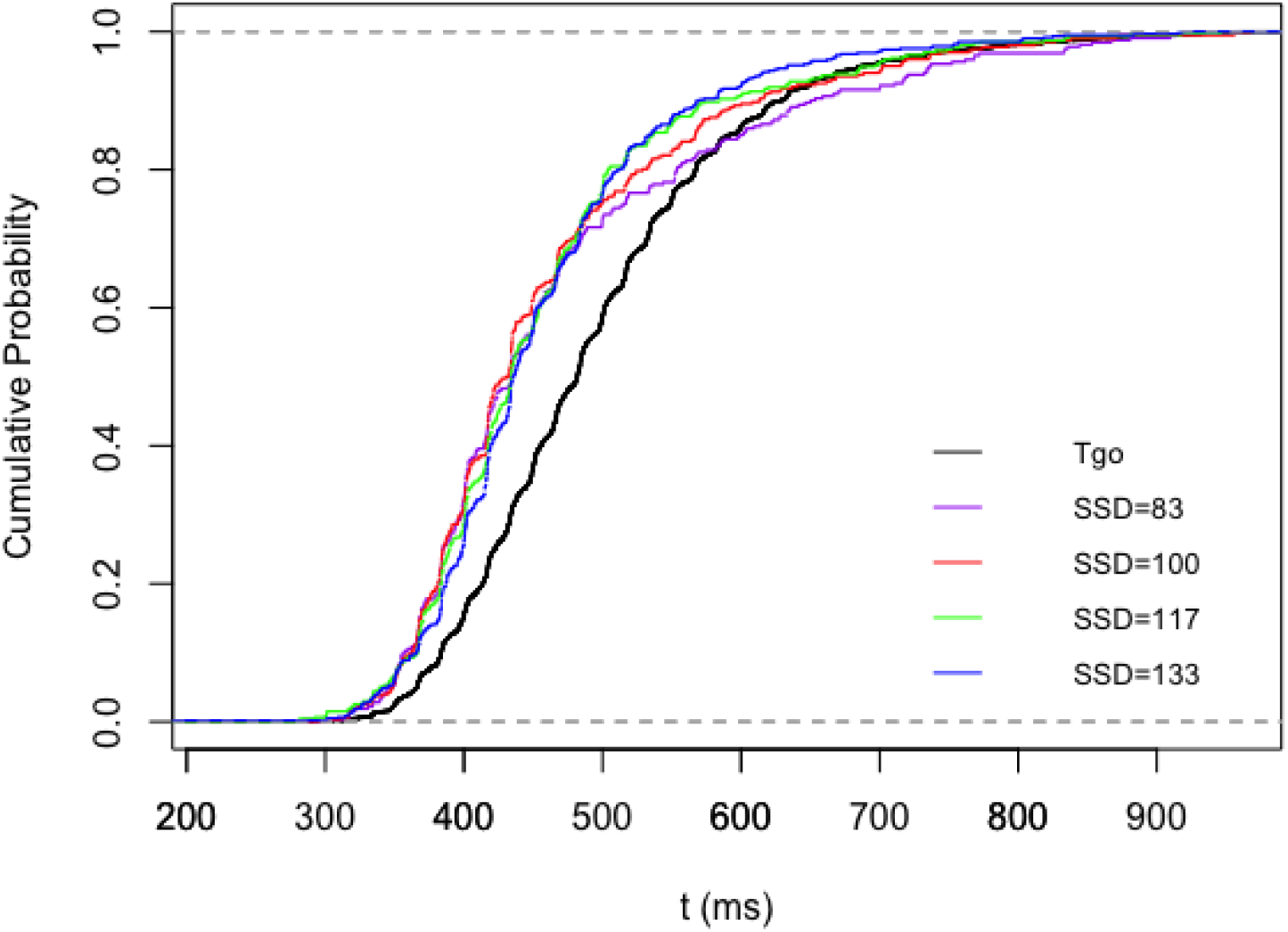
Cumulative distribution functions (CDFs) of response times (RTs) for testing inequality (2). The go distribution (*P* (*T*_*go*_ *< t*)) is shown with a black line, and the signal-respond distributions (*P* (*T*_*go*_ *< t* | *T*_*go*_ *< T*_*stop*_ + *t*_*s*_)) are shown for *t*_*s*_ = 83, 100, 117, 133 ms in purple, red, green, and blue, respectively.

### Test of the mixture hypothesis

We used the test developed by van Maanen and colleagues (van Maanen et al., 2016, 2014) (see also Couto et al., 2024). It assesses the evidence supporting the null hypothesis (i.e., the existence of the fixed-point property) versus evidence against it by quantifying the probability of observing the property compared to not observing it^5^. Both ANOVA and Bayesian ANOVA using the “*fixedpointproperty*” package in *R* were performed on the data, yielding *BF*_01_ = 0.34. This indicates that the data are 2.9 times more likely to demonstrate the fixed-point property than to not demonstrate it (see Figure 2). Moreover, the p-value of ANOVA (*P* = 0.192) is consistent with this result. Summing up, this analysis of the Bompas et al. (2025) data has generated support for the presence of a mixture of responses to the go and the ignore signal in context *ℐ𝒢𝒩𝒪ℛℰ*. However, non-observability of the distribution of ignore signals, *F*_*ig*_(*t*), precludes estimating the mixture parameter as a function of ISD, *π*(*t*_*i*_) and, thus, testing of the *late-ignore-signal hypothesis*.

**Fig. 2:**
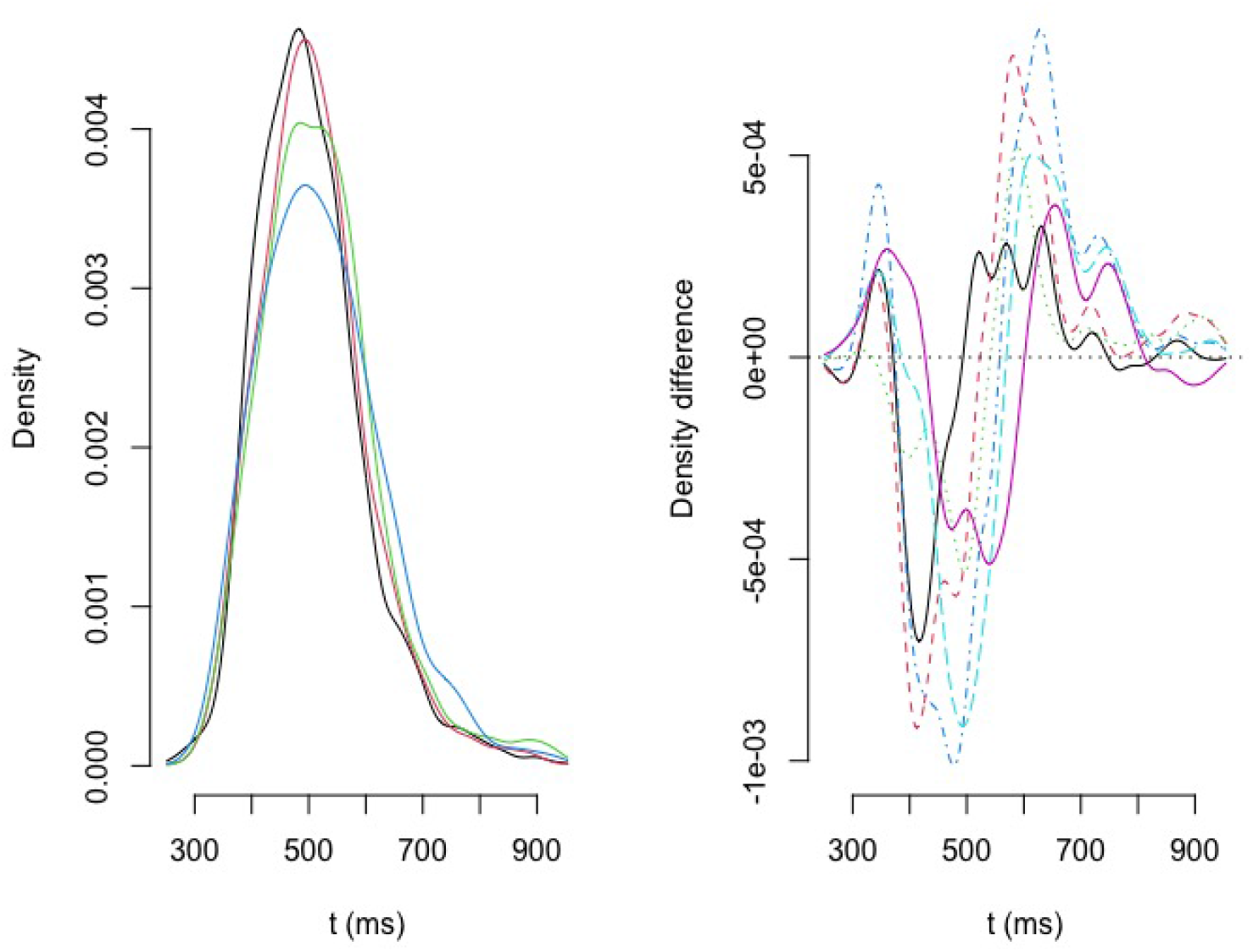
Plots of the 4 density estimates (left panel) and their differences (right panel) corresponding to ISD values *t*_*i*_ = 83, 100, 117, 133. At least one intersection point appears to lie between 300 and 400 ms, another one between 800 and 900 ms.

### Parameter estimation for the mixture model with shift (MMS)

The data from Bompas et al. (2025) feature 4 ISDs (83, 100, 117, or 133 ms) that are identical to the SSDs; we illustrate parameter estimation for the 4 mixture probabilities and the shift parameter *α*. After rewriting, we have 4 model equations for the means:

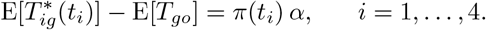

and 4 model equations involving the second raw moment (see Appendix):

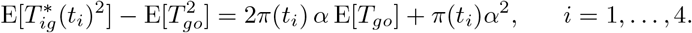

Inserting sample estimates for 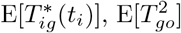, and 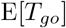, we end up with a system of 8 equations with 5 unknowns. Estimates for *π*(*t*_*i*_), *i* = 1, 2, 3, 4 and *α* are obtained by simultaneously minimizing the sum of the squared differences between the left- and right-hand sides of the equations (MatLab^©^ routine fminsearchbd). Using different starting values for the optimization (for *π*(*t*_*i*_) increasing, decreasing, or constant as a function of *t*_*i*_) yields different parameter values, but the estimate for *α* is always between 30 and 40 ms and the mixture probability estimates are always increasing with the ISD (*t*_*i*_) values, even if the starting values are set to decrease. The best sets of optimal values are listed in Table 1:

**Table 1:** MMS model parameter estimates for the data from Bompas et al. (2025) in MatLab^©^ routine fminsearchbd; SSD = (minimized) sum of squared deviations from model equations.

| $1 - \pi(83)$ | $1 - \pi(100)$ | $1 - \pi(117)$ | $1 - \pi(133)$ | $\alpha[ms]$ | SSD |
| --- | --- | --- | --- | --- | --- |
| .22 | .42 | .79 | .85 | 34.1 | 7.62 |

**Table 2:** Parameter estimates for the data from Bompas et al. (2025) for QMS model using the optim function (stats package) in the R programming environment.

| $1 - \pi'(83)$ | $1 - \pi'(100)$ | $1 - \pi'(117)$ | $1 - \pi'(133)$ | $\alpha'[ms]$ |
| --- | --- | --- | --- | --- |
| .36 | .51 | .79 | .85 | 34.7 |

Ignore signal delay *α* was found to be around 30 ms and estimates for the mixture probability are increasing in ISD. The estimates suggest that increasing ISD from 83 ms to 100 ms nearly doubles the likelihood that the response is triggered by the go signal rather than the ignore signal. We conclude that estimation of parameters for the mixture model with shift (MMS) is feasible and the dataset supports the late-ignore-signal hypothesis.

### Parameter estimation for the quantile-mixture model with shift (QMS)

After rewriting^6^, the equations for the quantile-shift model are, for any quantile level,

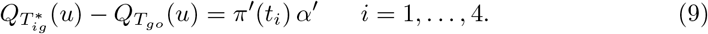

for the 4 ISDs (83, 100, 117, or 133 ms) in Bompas et al. (2025). With 5 unknown parameters, we need at least 2 different quantile levels in Equation 9 for estimation. Here, we chose 10 quantile levels, going from .05 up to .95 in steps of .05. Note that, compared to the mixture model with shift, it is much more straightforward to set up a large number of model equations for parameter estimation.

Similar to the procedure in the previous model, parameters were determined by simultaneously minimizing the sum of the squared differences between the left- and right-hand sides of the 40 equations of type (9), this time using the optim function (stats package) in the R programming environment.

These estimates are remarkably close, if not identical, to those of the mixture model with shift (MMS). This suggests that, for these data, it does not matter much whether the mixture occurs at the level of quantiles or distribution functions. Moreover, this finding may be interpreted to strengthen both the late-ignore-signal hypothesis and the general mixture assumption.

## Discussion

Stimulus-selective stopping is a variation of the standard stop-signal task in which, instead of a stop signal, an occasional ignore signal instructs participants to continue responding to the go signal. Several alternative strategies to perform successfully in this task have been proposed, and experimentally tested, in the literature (Bissett & Logan, 2014; Sanchez-Carmona et al., 2021). To our knowledge, no efforts have yet been made to develop a quantitative, non-parametric extension of the classic Logan-Cowan race model (Logan & Cowan, 1984) for this task. Here we introduce formal model classes that can quantitatively predict all observable data in the stimulus-selective paradigm.

We differentiate among three distinct contexts, depending on the type of signal presented in a given trial: contexts *𝒢𝒪, 𝒮𝒯𝒪𝒫*, or *ℐ𝒢𝒩𝒪ℛℰ* with corresponding random processing times for the go signal, the stop signal, and the ignore signal, denoted as *T*_*go*_, *T*_*stop*_, and *T*_*ig*_, respectively. In keeping with the classic race model, we propose that go and stop signal processing in contexts *𝒢𝒪* and *𝒮𝒯𝒪𝒫* follow the classic race model with stochastic independence between *T*_*go*_ and *T*_*stop*_, so that the usual predictions apply for this part of the data.

A naive extension of the classic model that would posit a race between *T*_*go*_ and *T*_*ig*_ on trials presenting an ignore signal (the *minimum model*), can be readily dismissed, as such a model would predict –via statistical facilitation (Raab, 1962)– faster responses in context *ℐ𝒢𝒩𝒪ℛℰ* – contrary to the commonly observed slowing.

The central and novel assumption of the models proposed here is that responses in the *ℐ𝒢𝒩𝒪ℛℰ* context are generated by sampling from a binary mixture of the distributions of *T*_*go*_ and *T*_*ig*_, i.e. *F*_*go*_ and *F*_*ig*_. This approach presents two main challenges. First, in contrast to *F*_*go*_, distribution *F*_*ig*_ is not observable, since a response in context *ℐ𝒢𝒩𝒪ℛℰ* could have been triggered either by the go or the ignore signal. Second, the mixture probability *π*(*t*_*i*_), of responding to the ignore signal presented with an ignore signal delay(ISD) of *t*_*i*_ [ms], is unknown as well. Fortunately, non-observability of *F*_*ig*_ did not prevent us from testing the mixture hypothesis, due to a test developed by van Maanen and colleagues (van Maanen et al., 2016) based on an earlier result in Falmagne (1968).

Note that the later the ignore signal occurs, the more likely it is that a response has been triggered by the go signal. This late-ignore-signal hypothesis implies that *π*(*t*_*i*_) is decreasing as function of *t*_*i*_. However, given the non-observability of the ignore signal distribution *F*_*ig*_, this hypothesis is not testable without being able to estimate the *π*(*t*_*i*_) values. This led us to introduce semi-parametric models that assume that the ignore signal processing distribution equals that of the go signal except for a constant delay, *α*. This, arguably, is a strong postulate, but it can be probed empirically in various ways.

We considered two classes of semi-parametric models that differ in the way the delay parameter is introduced in the mixture. In the first, called MMS, responses in context *ℐ𝒢𝒩𝒪ℛℰ* follow a mixture of distributions, as stated in Equation (4). In the second, called QMS, responses in context *ℐ𝒢𝒩𝒪ℛℰ* are defined as coming from a mixture of quantile functions, as stated in Equation (8). We developed methods to estimate all parameters, *π*(*t*_*i*_) and *α*, for both models. Although both model types postulate a mixture mechanism, their predictions may differ because, in general, the distribution corresponding to a mixture of quantile functions is not the same as distribution corresponding to a mixture of distributions.

Taking data from the study of Bompas and colleagues (Bompas et al., 2025), the test for the presence of a binary mixture (van Maanen et al., 2016) resulted in strong (Bayesian) support for the hypothesis. Furthermore, we used that dataset to illustrate the parameter estimation of the MMS and QMS models. We found plausible parameter values for both models with mixture probability estimates that are in line with the late-ignore-signal hypothesis. Estimates for the delay parameter were around 34 ms in both models. Although these findings support the strong assumption that the ‘ignore’ and ‘go’ distributions differ solely by a shift, a rigorous evaluation and comparison of the competing models has yet to be conducted.

In contrast to the mixture hypothesis, an alternative to generating a delay in the *ℐ𝒢𝒩𝒪ℛℰ* context is the introduction of a deadline. In early work, Ollman (1973) suggested that participants initiate a subjective “deadline” upon detection of the go signal and then either make a response if the deadline has terminated or withhold the response if the stop signal has been detected before deadline termination. In this variation of the classic race model, a response is observed in the *𝒮𝒯𝒪𝒫* context whenever the event

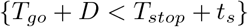

occurs, where *D* is a non-negative random variable representing the deadline. The distribution of *D* depends on the strategic decision of the participant: if avoiding inhibition errors is stressed, then *D* should be large enough to avoid the above event; if speed is stressed, then *D* should be reduced accordingly. Because participants do not know whether a stop signal comes up next, they should start the deadline process in the *𝒢𝒪* context as well. This model easily extends to comprise the *ℐ𝒢𝒩𝒪ℛℰ* context as well: a response should be observed whenever

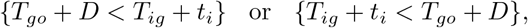

implying that responses occur with probability one again, if trigger failures are ignored. One issue with the deadline model is that the distribution of *D* is not observable, in addition to the distributions of *T*_*stop*_ and *T*_*ig*_. Moreover, introducing *D* as an additive component requires computing the convolution of distributions.

## Acknowledgment

This work was supported by DFG (German Science Foundation) grants to H. Colonius (CO 94/8-1) and A. Diederich (DI 506/18-1), and by the DFG Research Training Group (RTG) 2783 Project ID 456732630.

## Appendix : Mixture models with shift

For simplicity, we drop the dependence on *t*_*i*_ in Equation (4) and rewrite it as

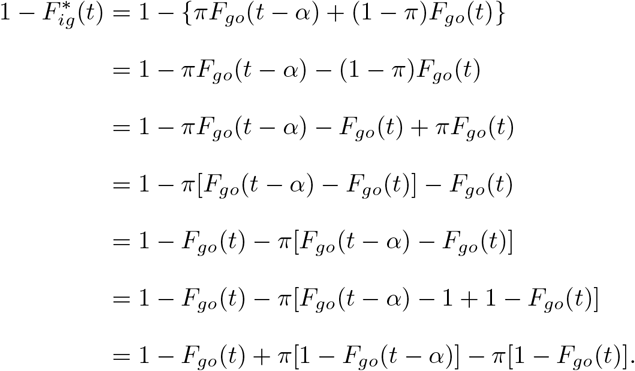

For the mean and *t >* 0, we take integrals:

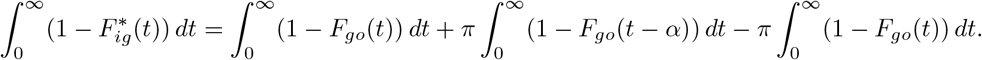

Yielding

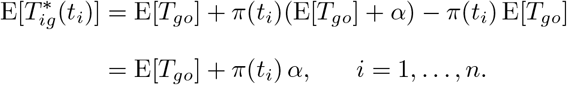

For the second raw moment, we first multiply both sides by 2*t*:

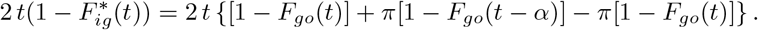

and integrate both sides (right side: summand per summand):

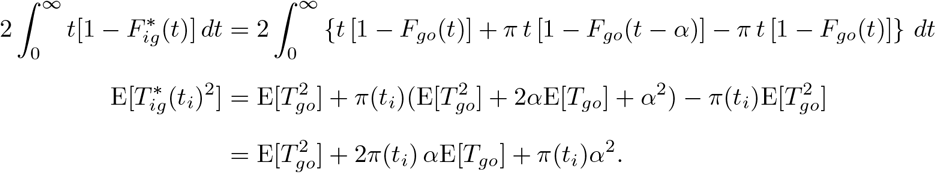

## Footnotes

1 aka as “context independence”

2 A further variant of stimulus-selective stopping involves a choice RT task with two different go signals; e.g., an arrow pointing to the left (or right) with instruction to press a left (or right) button (see Bissett & Logan, 2014).

3 We use subscript *ig, ℐ𝒢* as short for *ignore, ℐ𝒢𝒩𝒪ℛℰ*, respectively.

4 Equality only holds in special cases because the mixture distribution allocates different portions of probability to each component, which is not captured by averaging the quantiles.

5 Strictly speaking, the test does not apply here because one of the component distribution, *F*_*ig*_ (*t* − *t*_*i*_) depends on the mixture probability *π*(*t*_*i*_) by a shift. However, simulations in van Maanen et al. (2016) suggest that the effect is negligible compared to the one created by a change of the shape

6 We keep the primes to distinguish them from the parameters in the mixture model with shift.

## Notes

### Competing Interest Statement

The authors have declared no competing interest.

